# Microplastic pollution of commercial fishes from coastal and offshore waters, Japan

**DOI:** 10.1101/2021.10.20.465208

**Authors:** Mitsuharu Yagi, Tsunefumi Kobayashi, Yutaka Maruyama, Sota Hoshina, Satoshi Masumi, Itaru Aizawa, Jun Uchida, Tsukasa Kinoshita, Nobuhiro Yamawaki, Takashi Aoshima, Yasuhiro Morii, Kenichi Shimizu

## Abstract

Microplastics (MPs) pollution is a worldwide issue in the marine environment. There is growing concern of consuming MPs through fish, yet the current contamination status of fish collected from the deep sea surrounding Japan remains limited. We present baseline data on MPs in commercially important fishes from the coastal and offshore waters near Kyushu, Japan (East China Sea). We examined the MPs in the digestive tracts of two pelagic (n = 150 in total) and five demersal species (n = 235 in total). The fish were caught by pole and line, and bottom trawl at different geographical positions. The MPs in pelagic fish (39.1 %) were higher than demersal fish (10.3 %) and of larger sizes. The MPs correlated with habitat depth and type. There was species variation in the shape and polymer composition of MPs. These results increase our understanding of the heterogeneous uptake of MPs by fishes.

## 1. Introduction

There is growing global concern on microplastics (MPs) pollution in marine ecosystems (Klingelhöfer et al., 2020). Annual production of plastics has been increasing exponentially since the 1940s and 1950s, and now exceeds 350 million tonnes produced each year (Plastics Europe, 2020). Inappropriate management of plastic waste has caused contaminants to accumulate in the environment, resulting 20 million tons of plastic litter to enter the ocean every year (Vannela, 2012). Once plastic enters the ocean, it degrades and fragmentates into smaller pieces by ultraviolet radiation, oxidation, as well as wave and microbial degradation (Barnes et al., 2009; Cole et al., 2011; Moore, 2008). Plastics can also be crushed at a breaker zone (Isobe et al., 2015). The miniaturization of plastic (instead of decomposition) causes plastic particles smaller than 5 mm, these are defined as MPs (Barnes at al., 2009; Betts, 2008). MPs are tiny in size but degrade extremely slowly (Chamas et al., 2020), they drift in the marine environment for a long time and flow with currents throughout the borderless ocean.

The tiny particle size of MPs pollution of the marine ecosystem through their uptake by marine organisms. Ingestion of MPs occurs in a wide range of taxa such as invertebrates (e.g., Bour et al., 2018; Cho et al., 2019; Courtene-Jones et al., 2017), birds (e.g., Blight and Burger, 1997), turtles (e.g. Duncan et al., 2019), and fishes (e.g. Neves et al., 2015; Hyrenbach et al., 2020; Rasta et al., 2021; Wootton et al., 2021). MPs have been found in their gastrointestinal digestive tract of approximately 390 fish species (Savoca et al., 2021). Ingestion of MPs may cause physical and physiological damage (Wright et al., 2013). MPs accumulate persistent bio-accumulative and toxic chemicals (PBTs) as well as persistent organic pollutants (POPs) from the environment, and these toxins are transported into marine organisms through the uptake of MPs (Bakir et al., 2014; Rochman et al., 2013). Furthermore, the plastic additives contained in MPs can cause adverse effects (Teuten et al., 2009; Bejgarn et al., 2015). These ingested anthropogenic contaminants are consumed by human through trophic transfer (Carbery et al., 2018). Therefore, there is a need to clarify the status of these tiny contaminants in fish.

What type and where do fish ingest MPs? This critical information will increase our knowledge on microplastic pollution in fishes. The occurrence and frequency of the ingested MPs by fishes are thought to be influenced by the habitat (Jabeen et al., 2017; Baalkhuyur et al., 2018), trophic transfer (Bellas et al., 2016; Nikki et al., 2021) and feeding strategy (Romeo et al., 2015; Talley et al., 2020; Wootton et al., 2021). Low density MPs are abundant near the water surface (Suaria et al., 2016; Ory et al., 2017) and as density increases, the MPs decrease with increasing depth (Kukulka et al., 2012; Gorokhova, 2015). Isobe et al. (2015) reported that MPs density in the waters around Japan is 27 times higher than the world average. Therefore, we hypothesized the MPs incidence may be influenced by water depth as well as habitat type, i.e., pelagic versus demersal fish, and fishes residing in habitats near Japan (the East China Sea) have a high ingestion rate of MPs. In addition, there is limited information on the incidence of MPs in fish caught by commercial fishes in the waters around Japan (but see Tanaka and Takada, 2016), and virtually no research on the deep-sea area. To address these hypotheses, we conducted a MPs survey of the gastrointestinal tract (GIT) of fishes caught by pole and line, and the bottom trawl fishery, operating from 10 m to 167 m depths. Our study provides baseline data on the status of MPs uptake by commercial fishes and improves our understanding of the mechanisms aiding contamination by comparing various fish species and habitat types.

## 2. Materials and methods

### 2.1. Fish sampling

Fishes inhabiting the shallow coastal water along the west coast of Kyushu, Japan were sampled during three cruises at four station (St. 1 ∼ 4) from December 2017 to October 2018 using the training vessel *T/V* Kakuyo-maru (155 gross tonnage: Faculty of Fisheries, Nagasaki University) (Fig. 1 and Table 1). Fishes were categorized into pelagic, demersal and benthic species based on their habitat preference and behaviour (Friedman et al., 2020). In this study, the fishes were principally classified following Froese and Pauly (2021). The benthopelagic species inhabiting the near-bottom environment in the deep-sea (Marshall and Merrett 1977) were classified as demersal fish. Two pelagic fish species i.e., the chub mackerel *Scomber japonicus* (St. 1 and 2; n = 64 in total) and the Japanese jack mackerel *Trachurus japonicus* (St. 3 and 4, n = 40 in total) were captured by experimental pole and lines using underwater lamps (UL-3, Takuyo Riken Co. Ltd, Japan) at night after vessel had anchored at the station. Water depth at the fishing grounds ranged from 10 to 72 m (Table 1). However, pelagic fishes aggregate around lights at night (Inoue 1963; Lee at al., 2019) in fishing depths shallower than the anchored depth.

**Table 1.**
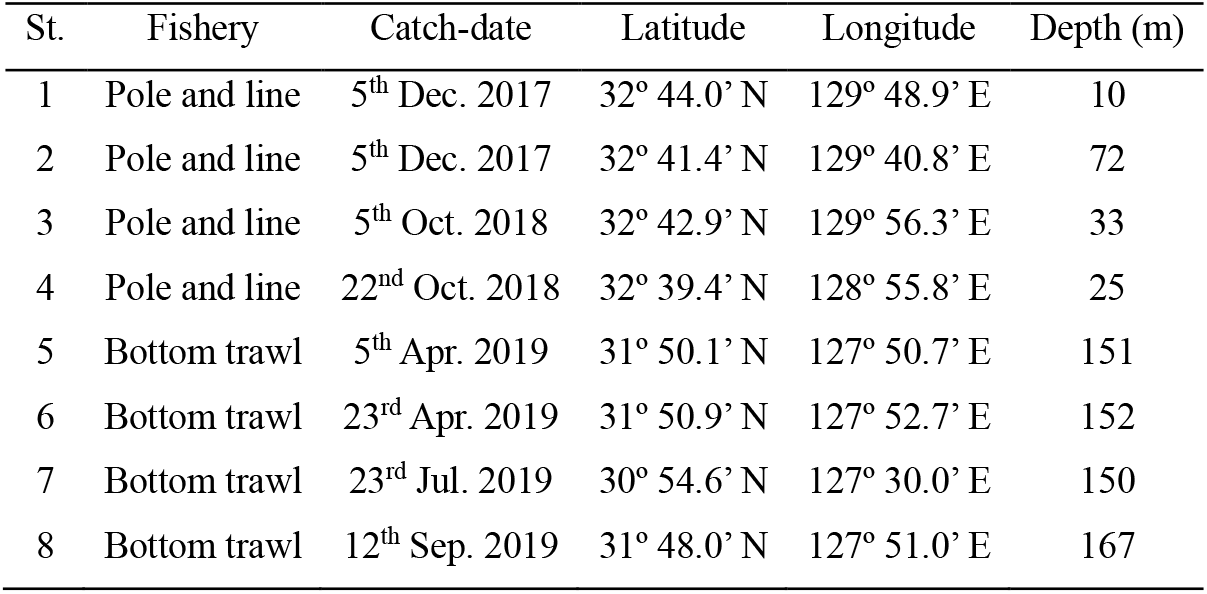
Summary of the fishery operation, catch-date, coordinates and water depth at each sample station in the East China Sea.

**Fig. 1.**
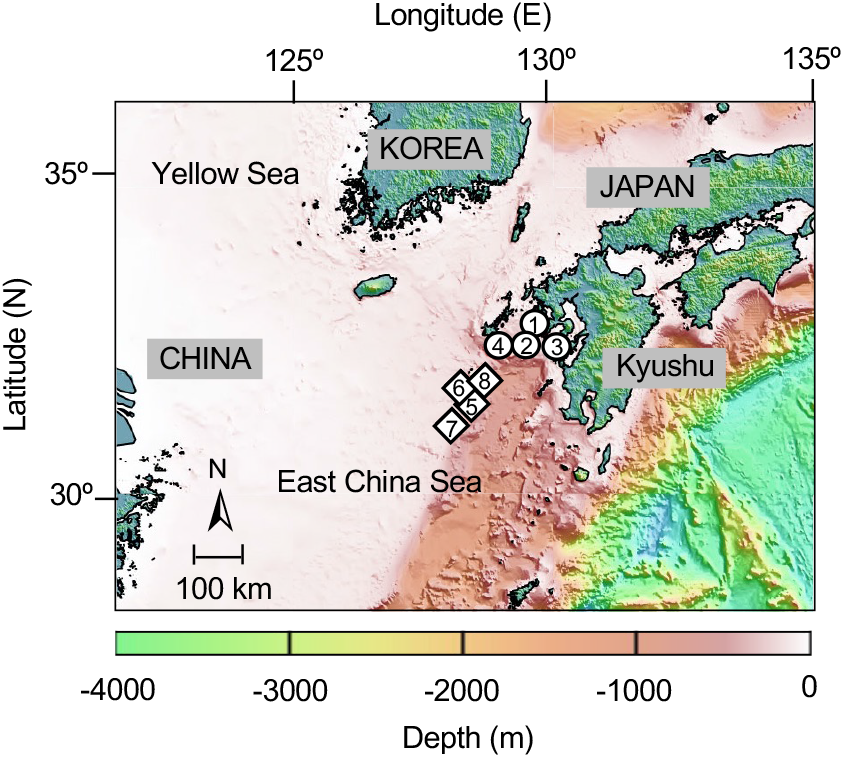
Bathymetric map demonstrating the geographical position of the sampling station in the East China Sea. The circles (St. 1 ∼ 4) and diamonds (St. 5 ∼ 8) signify the experimental pole and line and bottom trawl fishery, respectively.

An experimental bottom trawl was operated during four cruises at four station (St. 5 ∼ 8) from April to September 2019 in order to collect a greater diversity of fishes inhabiting deeper water using the *T/V* Nagasaki-maru (1,131 tonnage: Faculty of Fisheries, Nagasaki University). Five demersal fishes i.e., the yellowback seabream *Dentex tumifrons* (St. 5 and 6, n = 20 in total), the john dory *Zeus faber* (St. 5 ∼ 8, n = 59 in total), the longspine snipefish *Macroramphosus scolopax* (St. 5, n = 39), the redwing searobin *Lepidotrigla microptera* (St. 6, n = 44), the whitefin trevally *Carangoides equula* (St. 8, n = 73) and one pelagic fish, the Japanese jack mackerel *Trachurus japonicus* (St. 8, n = 46) were captured (Fig, 1 and Table 1). The trawler operated at water depths from 150 to 167 m. The trawl occurred during the daytime. A bottom trawl net with a 66 mm cod end mesh was towed for 30 minutes at approximately 2.0 knots. The initiation of the trawl time began when the net was set on the bottom and ceased when the net was retrieved off the bottom. All trawls were monitored acoustically. The size of the mouth opening and height of the trawl net (with the otter board) were approximately 24 m and 5 m, respectively.

All individuals were immediately stored at –20 °C in sealed plastic bags to prevent contamination before dissecting (Barboza et al., 2020).

### 2.2. Plastic sorting

The fish were dissected and the MPs were sorted in the Fish and Ships Laboratory, Faculty of Fisheries, Nagasaki University, Japan. To minimize potential contamination, standard provisions were undertaken (Kobayashi et al., 2021; Rasta et al., 2021). Briefly, we wore powder free Latex gloves and cotton lab. coats throughout the analysis process. All experimental equipment (including the desk) were rinsed with filtered, deionized water and cleaned with a 90% alcohol solution before and after each analysis. On completion of the assessment, all samples and equipment was covered with aluminium foil. We also utilised three petri dishes filled with ionized, filtered water to ensure the samples were free of any airborne contamination from the laboratory environment (Lam et al., 2020). A blank control was included in the analysis and was included in the dissection, digestion and microscopic observation and the visual examination. None of the blank controls contained any particles.

The MPs extraction procedure followed Rochman et al., 2015 with slight modifications. The characteristics of the surveyed fish species are presented in Table 2. After fishes were thawed at room temperature, the standard body length was measured to the nearest 1 mm using callipers. The wet weight was measured to the nearest 0.1 g using a digital balance. The fish were then dissected and the gastrointestinal tract (GIT) was removed. The GIT was weighed using a digital balance (with a 0.01 g accuracy). The GIT contents were carefully removed using fine tweezers. After removing the stomach contents, the GIT was weighed. The weight of stomach contents was calculated by subtracting the post GIT weight (without stomach contents) from the pre-GIT weight (with stomach contents), fish with small GIT were excluded from this analysis.

**Table 2.** Summary of the ingestion of microplastics by fish species sampled with pole and line, and bottom trawl fishery in the East China Sea. The size of MPs is provided in millimetres (mm).

| St. | Fish species | N.<br>(fish) | SL,<br>Mean ± S.D.<br>(max - min) | BW,<br>Mean ± S.D.<br>(max - min) | SC,<br>Mean ± S.D.<br>(max - min) | N.<br>(MPs) | MPs<br>incidence<br>(%) | Probable N.<br>of MPs per<br>individual | N. of MPs per fish with MPs<br>Mean ± S.D.<br>(max - min) | Size of MPs,<br>mean ± S.D.<br>(max - min) |
| --- | --- | --- | --- | --- | --- | --- | --- | --- | --- | --- |
| 1 | *S. japonicus*<sup>a</sup> | 40 | 152 ± 16<br>(182 - 105) | 50.4 ± 15.0<br>(73.0 - 19.6) | 1.58 ± 0.86<br>(3.41 - 0.04) | 38 | 42.5 | 0.95 | 2.24 ± 1.65<br>(8 - 1) | 1.95 ± 1.58<br>(8.20 - 0.46) |
| 2 | *S. japonicus*<sup>a</sup> | 24 | 289 ± 31<br>(355 - 228) | 243.9 ± 72.7<br>(434.4 - 112.0) | 5.91 ± 5.24<br>(27.73 - 1.36) | 47 | 62.5 | 1.96 | 3.13 ± 2.56<br>(9 - 1) | 4.83 ± 3.80<br>(17.34 - 0.30) |
| 3 | *T. japonicus*<sup>a</sup> | 38 | 177 ± 39<br>(228 - 112) | 102.5 ± 59.2<br>(210.2 - 19.2) | 1.77 ± 1.30<br>(7.32 - 0.53) | 17 | 31.6 | 0.45 | 1.41 ± 0.51<br>(2 - 1) | 2.47 ± 1.80<br>(7.34 - 0.50) |
| 4 | *T. japonicus*<sup>a</sup> | 2 | 117 ± 13<br>(126 - 108) | 31.6 ± 9.6<br>(38.4 ± 24.8) | NA | 1 | 50.0 | 0.50 | 1.00<br>(1 - 1) | 3.36<br>(3.36 - 3.36) |
| 5 | *D. tumifrons*<sup>b</sup> | 15 | 260 ± 21<br>(304 - 213) | 247.8 ± 72.1<br>(412.5 - 146.2) | 3.43 ± 1.43<br>(6.60 - 0.60) | 1 | 6.7 | 0.07 | 1.00<br>(1 - 1) | 8.80<br>(8.80 - 8.80) |
| 5 | *Z. faber*<sup>b</sup> | 9 | 165 ± 30<br>(220 - 128) | 150.0 ± 78.8<br>(333.8 - 60.4) | 2.90 ± 5.09<br>(16.34 - 0.21) | 3 | 22.2 | 0.33 | 1.50 ± 0.71<br>(2 - 1) | 1.75 ± 1.16<br>(3.00 - 0.70) |
| 5 | *M. scolopax*<sup>b</sup> | 39 | 161 ± 40<br>(227 - 97) | 25.1 ± 7.3<br>(35.3 - 8.7) | NA | 3 | 7.7 | 0.08 | 1.00<br>(1 - 1) | 1.50 ± 0.46 |
| 6 | *D. tumifrons*<sup>b</sup> | 5 | 225 ± 8<br>(231 - 214) | 426.1 ± 40.5<br>(484.0 - 383.8) | 5.18 ± 1.01<br>(6.47 - 3.67) | 0 | 0 | 0.00 | 0.00<br>(0 - 0) | NA |
| 6 | *Z. faber*<sup>b</sup> | 6 | 174 ± 11<br>(192 - 163) | 166.5 ± 29.7<br>(201.3 - 118.9) | 7.26 ± 5.71<br>(15.74 - 1.89) | 1 | 16.7 | 0.17 | 1.00<br>(1 - 1) | 0.40<br>(0.40 - 0.40) |
| 6 | *L. microptera* | 44 | 118 ± 9<br>(135 - 97) | 37.9 ± 8.5<br>(56.5 - 19.5) | NA | 0 | 0 | 0.00 | 0.00<br>(0 - 0) | NA |
| 7 | *Z. faber*<sup>b</sup> | 24 | 103 ± 16<br>(141 - 77) | 42.0 ± 21.0<br>(101.5 - 20.6) | 0.23 ± 0.11<br>(0.39 - 0.10) | 6 | 20.8 | 0.25 | 1.20 ± 0.53<br>(2 - 1) | 1.34 ± 0.53<br>(2.01 - 0.70) |
| 8 | *Z. faber*<sup>b</sup> | 20 | 110 ± 14<br>(141 - 91) | 46.8 ± 19.8<br>(107.0 - 29.2) | NA | 1 | 5.0 | 0.05 | 1.00<br>(1 - 1) | 1.33<br>(1.33 - 1.33) |
| 8 | *T. japonicus*<sup>a</sup> | 46 | 118 ± 9<br>(135 - 101) | 27.6 ± 5.4<br>(42.5 - 15.5) | NA | 6 | 8.7 | 0.13 | 1.50 ± 0.45<br>(2 - 1) | 2.90 ± 2.00<br>(6.72 - 0.96) |
| 8 | *C. equula*<sup>b</sup> | 73 | 108 ± 31<br>(184 - 77) | 61.3 ± 58.3<br>(283.3 - 16.4) | 0.40 ± 0.26<br>(0.88 - 0.05) | 11 | 13.7 | 0.15 | 1.10 ± 0.40<br>(2 - 1) | 1.80 ± 1.42<br>(5.77 - 0.61) |

Note: <sup>a</sup> pelagic fish, <sup>b</sup> demersal fish, N. = Number, SL = Standard length (mm), BW = Wet body weight (g), SC = weight of stomach content (g), MPs = Microplastics, and NA = Not available

The MPs collected from the stomach contents were cleaned using a chemical digestion method (Foekema et al., 2013; Karami et al., 2017). The stomach content was transferred to a jar, then 10 % KOH solution was added until the total volume reached 100 ml to dissolve the organic matter. The jar was incubated at 40 °C over 10 days (Tanaka and Takada, 2016). After chemical digestion, the contents were sieved through a 200 µm mesh sieve using filtered, deionized water. The sieved samples were sorted and examined under a dissecting microscope (SMZ745T, Nikon Corporation, Tokyo, Japan) using a glass petri dish. The sample debris were individually stored in pill cases (Kobayashi et al., 2021).

All particles were photographed using a digital camera (NOA630, WRAYCAM Corporation, Tokyo Japan) attached to a microscope. The longest section was measured from the image taken by the digital camera using *MicroStudio* software (WRAYCAM Corporation, Tokyo Japan). MPs were categorized according to their shapes (fragment, film and fibre) and colours (white, transparent, black, yellow, green, blue and others) based on the method described in Kobayashi et al. (2021) and Cheung et al. (2016). Finally, an attenuated total reflection Fourier transform infrared spectrometer (ATR-FTIR; FT-IR-4600, JASCO Corporation, Tokyo, Japan) was used to identify the plastic polymer composition of all the plastic samples, except the fibre shapes. The polymer type for identifying the fibre shape was not possible as they were below the minimum detectable quantity. Thus, we expressed the polymer type for the fibre shape as ‘fibre’. We applied ATR-FTIR to all cleaned MPs, which exhibited a spectrum range from 4000 to 400 cm^-1^ with a resolution of 4 cm^-1^. Eight scans were run for each measurement, and blank background scans were conducted every 50 samples. The diamond prism was cleaned with 75% ethanol prior to operating the ATR. The polymer type was identified by comparing with the reference libraries from KnowItAll Spectroscopy (Wiley Science Solutions, New Jersey, USA). The Hit Quality Index (HQI) between the tested sample and the reference library was recorded.

### 2.3. Statistical analyses

The significant differences in the MPs within different fish samples, the number of MPs, and water depth were identified using Pearson’s correlation tests. Pearson’s correlation tests were also analysed the relationships between the number of MPs, stomach contents and body weight. Data on the MPs incidence, and the number and size of MPs were tested using Bartlett’s test to confirm the equality of the variances. If the variances were unequal, alternative non-parametric tests were adopted (Zar, 2010). The variance analysis used the non-parametric Wilcoxon test or the parametric two-tailed Student’s t-test was performed. The non-parametric Kruskal Wallis test investigated multiple components among fish species within the same habitat type group i.e., demersal fish group. A Fisher’s exact test clarified any differences in presence/absence data for MPs incidence, shape, colour, and polymer type among fish species within the same habitat type group and among habitat types. *D. tumifrons* ingested only one MP item, therefore this species was excluded from the variance analysis. All statistical analyses were performed using JMP Pro 15. A statistical probability of *p* < 0.05 was considered significant.

## 3. Results

Six of the seven fish species sampled contained MPs (Table 2). Overall the MPs incidence and the average number of MPs per individual was 20.6 ± 19.4% (mean ± S.D.) and 0.36 ± 0.53 items (mean ± S.D). The MPs incidence and the average number of MPs per pelagic individuals was 39.1 ± 20.4% (mean ± S.D.) and 0.80 ± 0.71 items (mean ± S.D.) and per demersal individuals was 10.3 ± 8.40% (mean ± S.D.) and 0.12 ± 0.11 items (mean ± S.D.), respectively. The highest MPs incidence and highest average number of MPs per individual was found in *S. japonicus* (62.5 %, 1.96 items). The lowest MPs incidence and average number of MPs per individual was found in *L. microptera* (0 %, 0 item) (Table 2). The MPs incidence was strongly negatively correlated with fishing ground depth (Fig. 2a: *r* = -0.82, *p* = 4.0 × 10^−4^), accompanied with habitat type of the fishes (Fig. 2b: *Z* = 2.40, *p* = 0.016). The average number of MPs per fish followed the same pattern as the incidence (*r* = -0.62, *p* = 0.017) as well as habitat type (*t* = 2.87, *p* = 0.014) (Table 2). Fisher’s exact test showed the presence/absence frequency of MPs in the GIT were significantly different among fish species within same habitat type group (pelagic fish, *p* = 2.0 × 10^−4^, *n* = 152; demersal fish, *p* = 0.035, *n* = 235), implying the degree of MPs incidence variated between fish species even though they existed in the same habitat type.

**Fig. 2.**
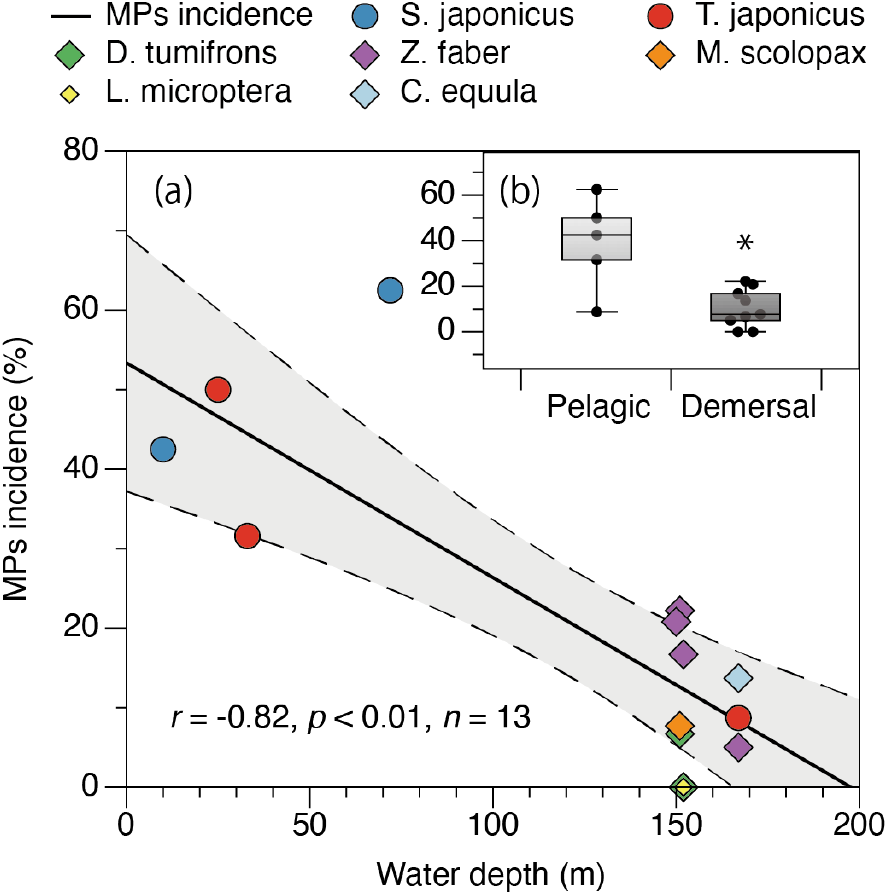
MPs incidence for fish species collected from different depths in Japanese waters. (a) Relationship between MPs incidence and water depth for all sampled species. The circles and diamonds signify pelagic and demersal fish species, respectively. The dashed line signifies the 95% confidence interval. (b) Comparison between fish with different habitat types using a whisker plot. The median (horizontal line), 25-75 percentiles (box) and range (bars) of the items ingested are presented. Statistically significant differences are indicated by * for *p* < 0.05.

The average number of MPs per fish was 1.88 ± 1.03 items (mean ± S.D.) and ranged from 9 to 0 depending on the fish species (Table 2). When further divided into fish type, the number of MPs per fish for pelagic and demersal fishes (mean ± S.D.) were 2.22 ± 1.52 items and 1.30 ± 0.35 items, respectively. Intraspecifically, the stomach contents positively correlate with body weight for *S. japonicus* (*r* = 0.65, *p* = 1.1 × 10^−8^), *T. japonicus* (*r* = 0.49, *p* = 0.002), *Z. faber* (*r* = 0.71, *p* = 1.0 × 10^− 4^) and *C. equula* (*r* = 0.80, *p* = 3.2 × 10^−6^), but not for *D. tumifrons* (*r* = 0.43, *p* = 0.056) (Fig. 3a). There was no relationship between the number of MPs per fish with MPs, the stomach contents and body weight, except for *S. japonicus*, which had an increase with stomach contents (*r* = 0.30, *p* = 0.021) (Fig. 3b) and with body weight (*r* = 0.26, *p* = 0.041) (Fig. 3c).

**Fig. 3.**
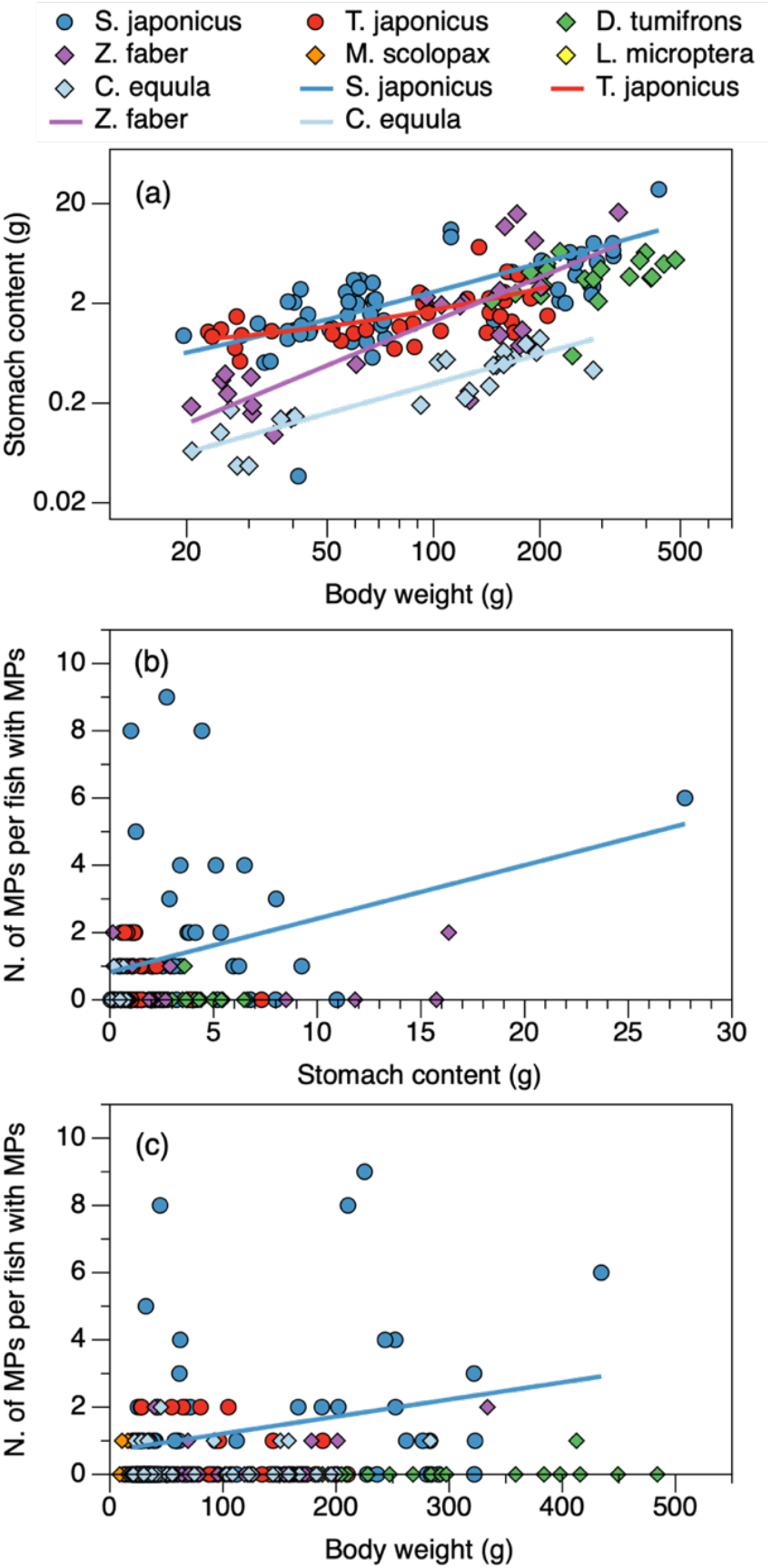
The relationship between number (N.) of MPs per fish with MPs, stomach content and body weight for all sampled species from coastal waters, Japan. The circles and diamonds signify pelagic and demersal fish species, respectively. Significant relationships (p < 0.05) are presented with regression lines. (a) Log stomach content vs log body weight. (b) N. of MPs per fish with MPs vs stomach contents. (c) N. of MPs per fish with MPs vs body weight.

The size of the ingested MPs was 3.05 ± 2.92 mm (mean ± S.D.), and ranged from 17.34 mm to 0.30 mm for all species. The mean size for the pelagic fishes and demersal fishes were 3.36 ± 3.12 mm and 2.05 ± 1.82 mm, respectively (Table 2). The mean size of MPs were not significantly different among fish species within the same habitat type group (pelagic, *χ*^*2*^ = 0.72, *p* = 0.397; demersal, *χ*^*2*^ = 5.4, *p* = 0.147) (Fig. 4a). The pelagic species ingested more larger MPs than the demersal species (*χ*^*2*^ = 9.63, *p* = 0.002) (Fig. 4b), implying the different habitat types affected the ingested MPs size. The quantification of the ingested MPs are summarized in Fig. 5. The HQI was 69.4 ± 16.3 (mean ± S.D.). The most dominant shapes of the MPs found in pelagic and demersal fishes was film and fibre, respectively (Fig. 5a). PE (polyethylene) and fibre were the dominant polymer type for pelagic and demersal fishes, respectively (Fig. 5b). Other polymers detected by ATR-FTIR were methyl methacrylate and Polyisoprene. The most common colours were transparent, white and blue, which comprised 70% of the MPs for both pelagic and demersal fish (Fig. 5c). Other colours included red, brown, and ash. One MP (n = 1) in a *D. tumifrons* was a fragmented, transparent and polyacrylonitrile. Fisher’s exact tests showed the shape and polymer types varied significantly between habitat types (shape, *p* = 1.4 × 10^−8^, *n* = 134; polymer type, *p* = 1.2 × 10^−8^, *n* = 134). Among fish species in the same habitat type group, there was not significant difference in shape of the MPs consumed (pelagic fish, *p* = 0.22, *n* = 109; demersal, *p* = 1.00, *n* = 25). In addition, the polymer type was significantly different between fish within same habitat group for pelagic fish (*p* = 2.4 × 10^−4^, *n* = 109), but the demersal fish showed significant differences between species (*p* = 0.48, *n* = 25). On the bases of colour, there was no significant difference between the habitat types (*p* = 0.09, *n* = 134) or among fish species within the same habitat groups (pelagic fish, *p* = 0.56, *n* = 109; demersal fish, *p* = 0.47, *n* = 25).

**Fig. 4.**
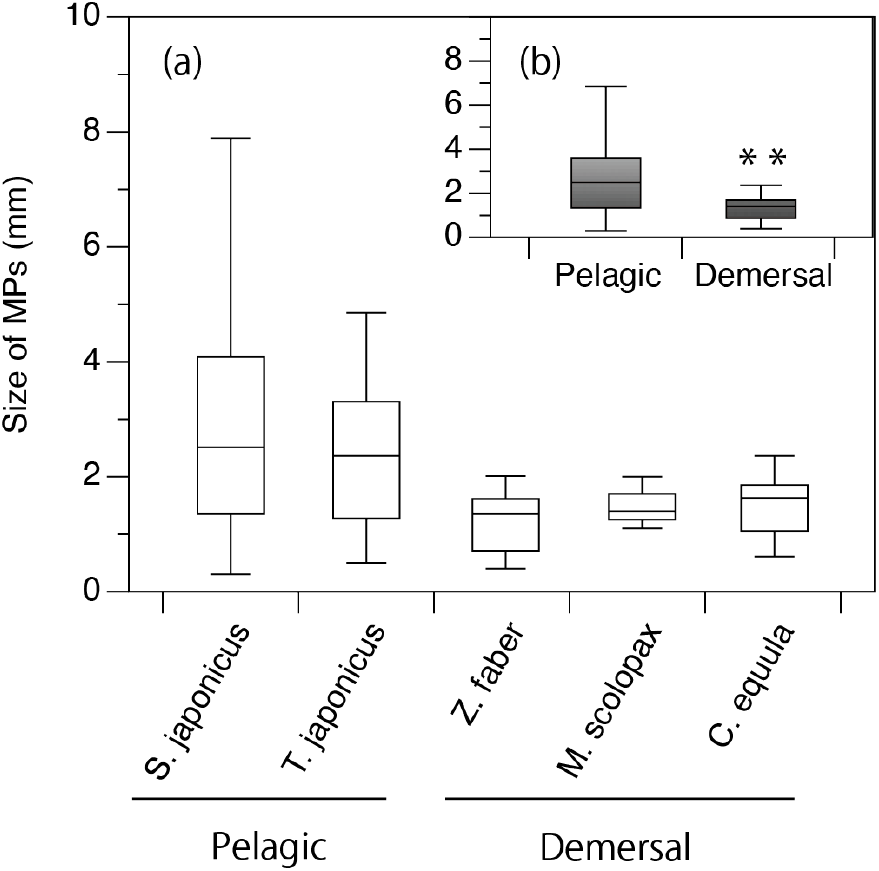
Size comparison of MPs in fishes from Japan. (a) Size of the MPs within each fish species. (b) Size of the MPs with fish habitat type. Presented are median (horizontal line), 25-75 percentiles (box) and range (bars) of the ingested MPs. Statistically significant differences are indicated by * * for *p* < 0.01.

**Fig. 5.**
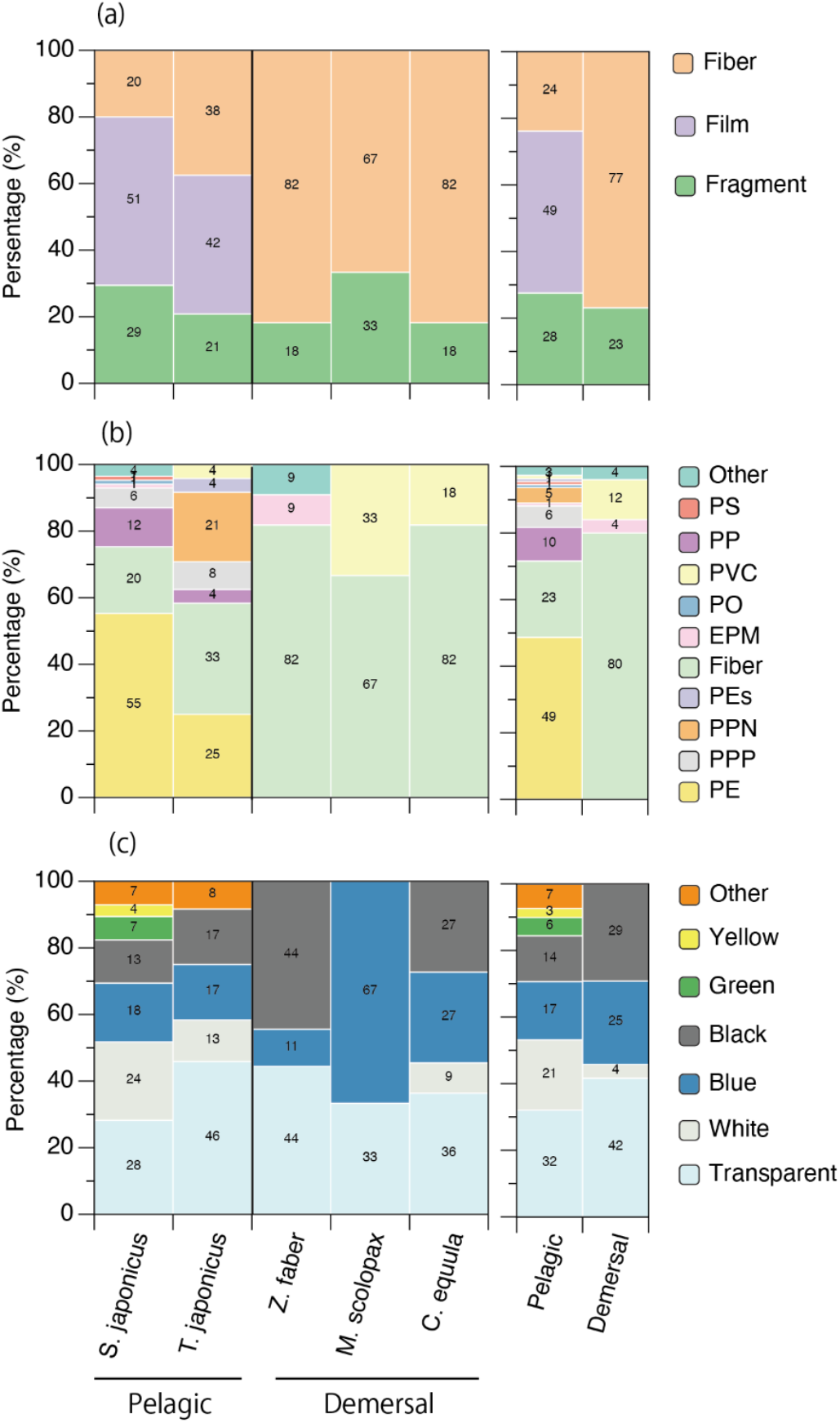
The composition percentage of MPs within fish species and habitat type of fishes from Japan. (a) shape, (b) polymer type and (c) MPs colour. EPM, ethylene propylene rubber; PE, polyethylene; PEs, polyester; PO, polyolefin; PP, polypropylene; PPN, poly-perinaphthalene; PPP, poly-pentaphenylene; PS, polystyrene; PVC, polyvinyl chloride.

## 4. Discussion

This is the first evaluation of the occurrence and abundance of MPs ingested by commercial fishes caught by bottom trawling the deep waters around Japan. As we hypothesised, the MPs incidence between fishes varied with the water depth of the fishing ground and the fish habitat type, i.e., pelagic vs demersal fish. The MPs incidence in pelagic fish (39.1 %) was statistically higher than that of demersal fish (10.3 %). Our result is consistent with previous reports on fishes in the North Sea and Baltic Sea (Rummel et al., 2016), but contrast with other studies (e.g., Lusher et al., 2013; Wootton et al., 2021). Pelagic fish are epipelagic feeders, and MPs are abundant near the water surface (Reisser et al., 2015; Suaria et al., 2016), whilst demersal fish are bottom feeders, and the abundance of MPs decreases with increasing water depth (Kukulka et al., 2012). An increase in the abundance of MPs leads to an increase in the bioavailability of MPs. Therefore, it is reasonable that pelagic fish ingest more MPs than demersal fish. However, detritivores demersal fish feed by extracting food from the sediment and are likely to have a higher frequency of ingestion (Jabeen et al., 2017; Wootton et al., 2021), because MPs accumulate on the sea floor (e.g., Barrett et al., 2020; Woodall et al., 2014). Interestingly, in the present study, pelagic fish *T. japonicus* captured from both the shallow and deep fishing grounds, showed the population in the deep water contained less MPs (Fig. 2). These results suggest the MPs abundance in the habitat is the most important factor in the ingestion of MPs by fishes.

The volume of MPs ingested by fishes is not only habitat-dependent, but also feeding strategy-dependent. We also hypothesized that most fishes in the waters around Japan would have a high rate of ingested MPs in the GIT because the area is a hotspot of sea surface MPs, it is 27 times higher than the world’s average (Isobe et al., 2015). In fact, Tanaka and Takada (2016) reported the planktivorous pelagic fish, the Japanese anchovy had ingested MPs at a high level (77%) in Tokyo Bay, Japan. However, in this study, the ingestion level in the pelagic fish was 39.1 %, which is close to the global average of 37.6 % (Markic et al., 2019). The reason for the lower level of MPs incidence may lie in the unique density distribution of MPs. Kobayashi et al. (2021) reported the difference between the highest and lowest abundances were 550-fold across all net tows, based on a survey conducted over a 14-month period near Kyushu, Japan. The spatiotemporal variations in MPs abundance affects MPs incidence in fish. Furthermore, the level of MPs incidence varied with fish species even though the species occurred in the same habitat type. This heterogeneity within the same habitat group of fish species may be due to the different feeding strategies. Among pelagic fish, *S. japonicus* had the highest MPs incidence. This is consistent with Neves et al. (2015) which examined 26 species of commercial fish off the Portuguese coast. Whereas *S. japonicus* is carnivorous, *T. japonicus* is a selective plankton feeder (Froese and Pauly, 2021). The opportunistic feeding strategy of *S. japonicus* may increase the MPs incidence. Interestingly, *L. microptera*, a benthic carnivorous demersal species (Togashi et al., 2019) ingested no MPs. The genus *Lepidotrigla* (including *L. microptera*) search for food (such as small crustaceans) using free pectoral rays (Finger, 2000). We speculate that fish with sensory feeding strategies with a taste-detecting organ could avoid consuming MPs.

Our results suggest many of the fish probably consumed MPs directly during normal feeding activities, mistaking MPs for small fish and crustaceans or inhaling it accidently (primary ingestion), not trophic transfer (secondary ingestion). Secondary ingestion occurs when prey that consumed MPs are eaten by a predator (Nelms et al., 2018). The mean number of MPs was 0.36 items per individual with MPs, which is nearly seven times lower than the global average of 2.6 items (Markic et al., 2019). The weight of the stomach contents increased with increasing body weight for four of the five species (Fig. 3a). If secondary ingestion of MPs occurred, the number of MPs will increase the food volume. However, the number of MPs per fish did not increase with the increasing weight of the stomach contents, except for *S. japonicus* (Fig. 3b). Considering the high variability in the number of MPs in *S. japonicus* specimens, the significant difference may be caused by outlying data. Thus, the MPs ingested by fish (including *S. japonicus*) in this study are considered primary ingested. We believe this is the first demonstration of the relationship between stomach contents and ingested number of MPs. The significant relationship between body weight and the number of MPs (Fig. 3c) may be caused by multicollinearity. Thus, further research must focus on the weight of the stomach contents instead of body size (e.g., Rahmawati and Patria, 2019; Lefebvre et al., 2019).

Pelagic fish may size-selectively ingest MPs. Pelagic fish (3.36 ± 3.12 mm) ingested more larger sized MPs than the demersal fish (2.05 ± 1.82 mm), and there was no difference among species within the same group (pelagic or demersal). In the same area, Kobayashi et al. (2021) reported the mean size of the surface MPs was 1.71 ± 0.93 (mm ± S.D., *n* = 6131) when the samples were collected throughout the year by neuston net with the a mesh size of 350 µm. Using this previous data and our study, there is a significant difference between the size of the surface MPs and the MPs found in fish GIT (Wilcoxon test, *Z* = 6.36, *p* = 2.0 × 10^−10^). Fish potentially select larger MPs when they feed on prey. Unfortunately, we cannot discuss the size-selectivity by demersal fish, because data is not available on the size distribution of MPs in the deeper water column in this area. Hence, more detailed studies are needed to examine the abundance and size distribution of MPs in the deeper water column and sediment.

There was qualitative differences in the ingested MPs within the fish species, i.e., the shape and polymer type among pelagic and demersal fish were characterised by film-shape, and low-density polyolefin. Pelagic fish mainly ingested fragments and films made from polyolefin (e.g., PE and PP). The polymer type identified in fish in our study support previous findings by Wootton et al., (2021), where fish in southern Australia mainly ingested polyolefin. Both PE and PP are buoyant polymers with a lower density than seawater (Lefebvre et al., 2019). Kobayashi et al. (2021) identified the frequency of polymers and the colour of the surface MPs in the same area closely resembled the composition of the polymer type of the MPs ingested by pelagic fish. In our study, the significant differences in the polymer type composition between *S. japonicus* and *T. japonicus* may have been influenced by the habitat depth. Unlike *S. japonicus, T. japonicus* occurs near the bottom layer during the daytime and were the dominant species collected in the bottom trawl survey in the East China Sea (Ohshimo, 2004; Sassa et al., 2009). The depth of the waters where *T. japonicus* resides may lead to less lower density polyolefin ingested (67 % for *S. japonicus* and 29 % for *T. japonicus*) (Fig. 5b). The demersal fish species ingested high density polymer PVC, which is heavier than seawater (Morét-Ferguson et al., 2010), with less collected in the surface samples (Kobayashi et al., 2021; Lefebvre et al., 2019). Thus, the qualitative composition of MPs ingested by fish may reflect the environmental composition of the MPs. Finally, the dominant polymer in demersal fish was fibre. This result is consistent with Murray and Cowie (2011), who surveyed bottom dwelling finfishes and shell fish in a blackish water lake in India. Mizraji et al. (2017) also showed a high abundance of fibres in the GIT of fishes in the upper tidal pools in Chile. Murray and Cowie (2011) identified fibre as particularly hazardous, because they can clump and knot, potentially preventing egestion. In a rearing experiment, Au et al. (2015) showed that during acute exposure, PP fibres were more toxic than PE particles. Fibres should be distinguished from other polymer types in both their distribution in the environment and impact on living organisms.

Fish, like humans are constantly ingesting MPs. In this study, the mean size of the MPs was approximately 3 mm, and egested by the fish. Batel et al. (2016) and Cedervall et al. (2012) reported the egesting time of MPs ranges from days to weeks. Although surveying MPs ingestion by fish provides only ‘snapshot’ data, the fish will be routinely consuming MPs. Fish species in this study are commercially important with the local human population frequently eat them. For example, *S. japonicus* which registered highest MPs incidence, is one of the most important commercial middle-sized pelagic fish, and a cosmopolitan species with a wide distribution throughout the Atlantic and Pacific and adjacent oceans (Collette and Nauen, 1983). Although people seldom consume the entire body of a fish (including GIT), some studies indicate MPs are also in the tissues of fish (Abbasi et al., 2018; Collard et al., 2017). Thus, humans are constantly consuming MPs within commercial fish. The potential impact on human health by the consumption of fishes is not well understood (Carbery et al., 2018; Vethaak and Legler., 2021). The abundance of MPs in the environment is linked to the presence of MPs in fish and their biota (Ferreira et al., 2020; Tien et al., 2020; Zhang et al., 2020). We need to aim for a “zero-plastic-ocean” as individuals, local businesses and worldwide.

## 5. Conclusion

This study is the first to document the presence of microplastics in the gastrointestinal tracts of two pelagic and five demersal fish species from coastal and offshore waters near Kyushu, Japan. Six of the seven fish species caught by pole and line, and bottom trawl fishing had microplastics present in their gastrointestinal tract. The MPs incidence and the number of MPs ingested varied with fishing ground depth, habitat type and feeding strategy of the fish species. This study provides an important contribution to our knowledge and understanding of plastic pollution in these commercial fish.

## Acknowledgements

We are grateful to all the crew on the training ships (*T/S*) *Nagasaki-maru* and *Kakuyo-maru* for their support and safe ship navigation. We also acknowledge the “Fish and Ships Laboratory” students from the Faculty of Fisheries, Nagasaki University, who assisted us. Finally, thanks to the editors and anonymous reviewers for their valuable comments and suggestions that greatly improved the quality of the manuscript.

## CRediT authorship contribution statement

**Mitsuharu Yagi**: Conceptualization, Methodology, Investigation, Writing – original draft, Supervision, Funding acquisition, Visualization. **Tsunefumi Kobayashi**: Investigation, Writing – review & editing. **Yutaka Maruyama**: Investigation, Writing – review & editing. **Sota Hoshina**: Investigation, Writing – review & editing. **Satoshi Masumi**: Investigation, Writing – review & editing. **Itaru Aizawa**: Investigation, Writing – review & editing. **Jun Uchida**: Investigation, Writing – review & editing. **Tsukasa Kinoshita**: Investigation, Writing – review & editing. **Nobuhiro Yamawaki**: Investigation, Writing – review & editing. **Takashi Aoshima**: Investigation, Writing – review & editing. **Yasuhiro Morii**: Investigation, Writing – review & editing. **Kenichi Shimizu**: Investigation, Writing – review & editing.

